# “Frustratingly easy” domain adaptation for cross-species transcription factor binding prediction

**DOI:** 10.1101/2025.05.21.655414

**Authors:** Mark Maher Ebeid, Ali Tuğrul Balcı, Maria Chikina, Panayiotis V Benos, Dennis Kostka

## Abstract

**Motivation:** Understanding how DNA sequence encodes gene regulation remains a central challenge in genomics. While deep learning models can predict regulatory activity from sequence with high accuracy, their generalizability across species—and thus their ability to capture fundamental biological principles—remains limited. Cross-species prediction provides a powerful test of model robustness and offers a window into conserved regulatory logic, but effectively bridging species-specific genomic differences remains a major barrier.

**Results:** We present MORALE, a novel and scalable domain adaptation framework that significantly advances cross-species prediction of transcription factor (TF) binding. By aligning statistical moments of sequence embeddings across species, MORALE enables deep learning models to learn species-invariant regulatory features without requiring adversarial training or complex architectures. Applied to multi-species TF ChIP-seq datasets, MORALE achieves state-of-the-art performance—outperforming both baseline and adversarial approaches across all TFs—while preserving model interpretability and recovering canonical motifs with greater precision. In the five-species transfer setting, MORALE not only improves human prediction accuracy beyond human-only training but also reveals regulatory features conserved across mammals. These results highlight the potential of simple yet powerful domain adaptation techniques to drive generalization and discovery in regulatory genomics. Crucially, MORALE is architecture-agnostic and can be seamlessly integrated into any embedding-based sequence model.

**Availability:** Code is available at https://github.com/loudrxiv/frustrating.

## 1 Introduction

Genomic regulatory activity is largely governed by transcription factors (TFs) that bind to DNA and influence gene expression. However, a fundamental question remains: can we accurately predict TF binding from sequence alone to determine regulatory function and cell identity? Sequence-to-function (S2F) models, typically implemented as deep neural networks, have become a cornerstone for addressing these questions and have facilitated a wide range of genomic analyses (e.g., downstream interpretation, prediction, functional element discovery, in silico sequence perturbation, and directed sequence design [1–3]). State-of-the-art models use convolutions, pooling, and autoregressive components to annotate sequences functionally and have been successfully applied to many tasks, revealing principles of gene regulation and guiding the analysis of genetic variations [4–7]. A crucial question, however, is the extent to which these models learn universal biochemical principles that allow generalization to out-of-distribution tasks.

An underexplored area of current machine learning techniques is moving beyond reliance on data from a single domain. Models exposed to data from only one domain are more likely to overfit to domain-specific features, which hampers generalization and weakens predictive power. Training on sequence data from multiple species, including model organisms with unique functional annotations, offers a promising approach to improving generalization. Specifically, training models with a multi-domain dataset could help identify underlying biological principles that are robust to species-specific variations while capturing shared, conserved information. However, can we assume that genomic elements with conserved regulatory functions between humans and model organisms are detectable through sequence alone? While some sequence-conserved regions can be identified based on multiple sequence alignments, at a finer scale, the majority of sites that bind transcription factors (TFs) are subject to rapid turnover — even between closely related species — making these sites difficult to annotate or characterize [8]. Yet, there is promise: the amino acid sequences of TFs, particularly their DNA-binding domains, are remarkably conserved, often showing high whole-protein sequence similarity across diverse species, alongside strong intrinsic DNA sequence preferences. This high degree of conservation in TF structure suggests a conserved “vocabulary” encoding rules of gene regulation. It also could suggest a cross-species gene regulatory grammar—a set of underlying rules governing TF-DNA interactions and their regulatory roles—that might be discovered using machine learning models trained on multi-species sequence data [9–12].

Recent methods have begun using sequence data from humans and model organisms to characterize regulatory elements, achieving some success. Kelley et al. [9] found that training Basenji (a convolutional model with pooling and dilated residuals) on combined human and mouse data increased test set accuracy on CAGE annotations (Cap Analysis of Gene Expression, a technique for mapping transcription start sites) compared to models trained on human data alone. Cochran et al. [13] found that when predicting transcription factor binding, a joint model (a CNN with an autoregressive component operating on 500-bp windows) trained on human and mouse data improved performance in predicting binding in humans. Their model used an adversarial approach (gradient reversal, GRL), training a classifier head to predict binding in the target species, alongside an adversarial discriminator that predicts the sequence’s species of origin. Gradients from this discriminator branch are penalized to encourage the model to learn species-invariant features. This approach requires both the parameterization of an adversarial branch.

Current approaches for cross species TF binding do not directly target latent embeddings to encourage or enforce an invariant representation of input sequences. One such approach is the the correction/alignment of moments from multiple domains of data [14]. Unlike the gradient reversal layer (GRL), which requires extra parameters for modeling domain discrimination, moment alignment can be expressed in closed form. This eliminates the need for additional parameters and allows for integration into any model with an embedding layer.

We offer MORALE, a framework that is “frustratingly easy” in that we implement a moment alignment scheme (we use first and second moments) to built on previous work [15]. We compare our method with existing ones [13] to predict the binding of two sets of transcription factors in humans and mice, and we perform a multi-species evaluation. Our framework shows strong overall predictive performance and learns a robust species-invariant feature set.

## 2 Materials & Methods

### 2.1 Data pre-processing

#### 2.1.1 Two-species case

We follow the procedure of Cochran et al. [13] to process ChIP-seq data from four TFs: CTCF, HNF4*α*, RXRA, and CEBPA in human and mouse liver tissue. ChIP-seq experiments and corresponding controls were collected from three sources: ENCODE, the NCBI Gene Expression Omnibus, and ArrayExpress, with the experiments and controls for each are listed under the following accession numbers: **(1)** ENCODE: ENCSR000CBU, ENCSR911GFJ, ENCSR098XMN, **(2)** ArrayExpress: E-TABM-722, and **(3)** Gene Expression Omnibus: GSM1299600.

Datasets were constructed into 500-bp windows, with a 50-bp overlap. We remove any windows marked as ENCODE blacklist regions [16]. FASTQ files were aligned to the human and mouse genomes (GRCh38, and GRCm38 respectively) using BowTie2 [17]. Peak-calling was performed using multiGPS v0.75 with default parameters [18]. Information on called peaks can be seen in **STab. 7**. Data was subsequently binarized for the 500-bp windows, and a window is called ‘bound’ if covers a peak’s center, and ‘unbound’ otherwise. In order to create balanced data for model training, we manually construct each minibatch of samples to contain an equal number of bound and unbound examples (for the source species, where this data is available). Because positive examples are sparse, they are shuffled and re-used more often than negative examples. For the domain-adaptive tasks, we also construct a ‘background’ set of random samples from each species, balanced based on species identity.

In order to construct validation and test sets for model evaluation, chromosomes 1 and 2 were held out from all training datasets. At the end of each training epoch, we evaluate 1 million randomly sampled windows from chromosome 1 and select the best-performing model. In order to assess model performance, we evaluate on all windows from chromosome 2. Sex chromosomes were excluded from training, validation and test sets.

#### 2.1.2 Multi-species case

We used ChIP-seq data in liver tissue across five species (human, rhesus macaque, mouse, rat, and dog) for four TFs (CEBPA, FOXA1, ONCECUT1 (HNF6), and HNF4*α*) from a previous study [19]. All files are available on Array Express, ID E-MTAB-1509. FASTQ files were aligned to the reference genome; we performed peak-calling using multiGPS v0.75 with default parameters. Filtered peaks from this analysis are in **STab. 8**. Filtering was done with SAMtools and Sambamba [20, 21]. We split the genomes into 1000-bp windows with a 50-bp overlap. Where applicable (i.e., mouse and human genomes) we removed any windows that are marked as ENCODE blacklist regions. Genomic data for processing (e.g., the references, BowTie2 indices, and blacklists) were obtained using the python package genomepy [22]. Binary window-labels were determined as in the two-species case (see **Section 2.1.1**).

Validation and test-sets were chosen differently from the two-species case, using an automatic approach to identify chromosomes approximating a pre-specified fraction of windows and covering a similar fraction of positive and negative windows, as we reported before [23]. Briefly, we solve the following optimization problem:

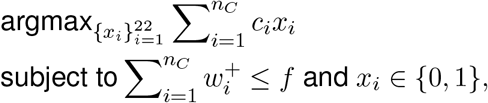

where *n*_*C*_ is the number of chromosomes, *f* is the desired fraction of positive windows, 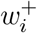 the fraction of positive windows on chromosome *i* (compared to all positive windows genome-wide), 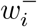 the corresponding fraction of negative windows, and 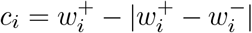. The optimization problem was solved using the R-packafge lpSolve. Selected chromosomes were held out for validation or testing, and excluded from training. Sex chromosomes were excluded. Similar to the two-species case, we re-use positive examples to create label-balanced minibatches shuffling and re-using positive windows more frequently than negative windows. We train over the course of 15 epochs and save the best performing model based on the target auPRC validation at the end of every epoch. For background sequences used for domain adaptive tasks, in each minibatch we randomly choose, without replacement, sequences from all chromosomes other than the validation, test, and sex chromosomes. Sequences are balanced by species identity. Overall batch construction is therefore similar as in the two-species case: a label-balanced part and a species-balanced (background) part that is label-agnostic but used for domain-adaptive losses.

### 2.2 Model Architecture

#### 2.2.1 Two-species case

We use the same network architecture as Cochran et al. [13] for this study. The network takes in 500-bp windows of one-hot encoded nucleotide sequences and passes it through an initial convolution layer with 240 20-bp-wide filters. This is followed by a pooling layer of size 15, an LSTM with 32 internal nodes, and then dense layers for the classification branch.

MORALE does not require a separate branch domain classification, so we simply calculate the moments based on the learned embedding after the LSTM (a vector of 32 internal nodes). We adapt the implementation from the python package ADAPT [24] in order to do so. For the gradient reversal, the instantiation of an adversarial discriminative branch is also defined. Following Cochran et al. [13], after max pooling this branch of the network is reshaped into a vector of size 8,160 and feeds into the GRL. The GRL merely outputs the identity of its input during the feed-forward step of model training, but during backpropagation, it multiplies the gradient of the loss by a factor of *™λ*. This layer is followed by a flatten layer, a ReLU-activated fully connected layer with 1024 neurons, a sigmoid-activated fully connected layer of 512 neurons, and, finally, a single-neuron layer with sigmoid activation. Notably this domain-classification branch increases the number of parameters of the overall network from 613,498 to 9,495,666. The domain-classification part, for GRL, therefore contributes the vast majority of model parameters.

#### 2.2.2 Multi-species case

We construct our model drawing from the gReLU collection of architectures [25], particularly adapting a model that contains a bidirectional GRU. The network consumes one-hot encoded windows of 1000-bp length, followed by 1-D convolutional layer with 240 filters of 20-bp width, and ReLU activation. Next, we apply a 1-D max-pooling layer of size 16. The output of the pooling is fed into single-layer bidirectional GRU with a hidden size of 240 for each direction. This is followed by two fully-connected layers with non-linear activations (ReLU). Next, we apply size-1 convolutions with 63 channels, followed by global average pooling. This forms the input for MORALE’s moment alignment [24], and for a linear classification head with sigmoid activation. In total, this network contains 960,367 parameters.

### 2.3 Model Training

#### 2.3.1 Two-species case

All models were trained using TensorFlow 2.18 [26, 27]. The Adam optimizer was used with a learning rate of 1e-3 for all models. Models were trained for 15 epochs. Following the protocol in Cochran et al. [13] for direct comparison, models were saved based on the best validation performance achieved on the source genome’s validation set. Batch size in the supervised setting was 400 (200 bound and 200 unbound examples from the source species). For domain adaptation, and additional 400 random regions were sampled for each species. We use the scikit-learn [28] implementation of average_precision_score to estimate the auPRC, which is our primary performance metric.

#### 2.3.2 Multi-species case

We train all models with PyTorch 2.5.1 [29] using the Adam optimizer with a learning rate of 1e-3. Models were trained for 15 epochs, with validation and test chromosomes constructed as discussed above (see **Section** 2.1.2). Minbatches were constructed to contain 200 bound and unbound examples, respectively, with equal contributions from all source species. The domain-adaptive task operates on 250 random examples (50 from each species, including the target). We validate all sources and the target species at the end of every epoch. The best performing model, on the target, during the course of training (based on validation performance, auPRC) is saved for test set evaluation. Like in the two-species case use `average_precision_score` in order to estimate auPRC for performance assessment.

### 2.4 Hyperparameter tuning

#### 2.4.1 Two-species case

For the GRL approach, to find *λ*, we construct a grid of values from 0.0 to 10.0 with a step size of 0.50 and select the maximal auPRC value for each transcription factor we study. We find the following: In the mouse-to-human direction (i.e., human is the target-species), we select 1.5, 6.0, 0.5, 7.5 for CTCF, CEBPA, HNF4*α*, and RXRA, respectively. In the human-to-mouse direction (i.e., mouse is the target species, we select 6.5, 8.5, 10.0, 1.0 for CTCF, CEBPA, HNF4*α*, and RXRA, respectively.

For MORALE’s hyperparameter (see **Equation 1**) we use a range from 0 to 10 with a step size of 1. In the mouse-to-human direction (i.e., human is the target-species), we select 4, 7, 8, and 8 for CTCF, CEBPA, HNF4*α*, and RXRA, respectively. In the human-to-mouse direction (i.e., mouse is the target species, we select 4, 8, 6, and 7 for CTCF, CEBPA, HNF4*α*, and RXRA, respectively.

#### 2.4.2 Multi-species case

We follow a similar procedure to tune MORALE in the multi-species case. We construct a tuning grid ranging from 1 to 8 with a step size of 1, for computational efficiency and based on the range of optimal values found in **Section 2.4.1**.

### 2.5 Differentially predicted site categorization

We follow Cochran et al. [13] in defining a “differentially predicted site” to quantify site enrichment between models. Specifically, we focus on differentially predicted false positives and false negatives to compare domain-adaptive models. Across five folds of data that we use to construct performance measures, we average model output for each window in our chromosome 2 test set, with either mouse or human as the target species. If the average sigmoid value for a window exceeds 0.5, we classify it as a “bound” prediction; if it is less, we classify it as “unbound.

To identify errors not made by the target-trained model but made by the source-trained model we focus on sites where the difference between the outputs of the source-trained model (no domain adaptation, DA) and the target-trained model exceeds 0.5. A differential false positive is a false positive from the source-trained model but a true negative in the target-trained model. A differential false negative is defined similarly.

### 2.6 Attribution & Importance scoring

For a sequence window and model, we generate both ‘attribution’ and ‘importance’ scores. Attribution scores estimate the per-nucleotide contribution as measured by expected integrated gradients. Specifically, we use the contribution_scores function within the Cis Regulatory Element Sequence Training, Explanation, and Design (CREsted, [30]) package. To create an overall importance score, we average over the channel dimension.

To capture over-represented sequence patterns to generate Contribution Weight Matrices (CWMs), we use the motif discovery tool TF-MoDISCo lite [31]. We focus on capturing the motif of the transcription factor (TF) that each model is trained on. Therefore, we subset the overall set of test windows to include only true positive sites sites where the model produced a sigmoid value of at least 0.98 across the five folds, defining a ‘strong’ true positive site. We randomly sample 2,000 of these sequences when applicable. For the parameters of TF-MoDISco, we use a maximum of 1,000,000 seqlets per cluster on a 500-bp window. In order to annotate found motifs and their corresponding CWMs, we use Tomtom [32] to label the patterns uncovered by TF-MoDISCo.

### 2.7 Repeat analyses

All analyses of repeat elements used RepeatMasker tracks for the corresponding genome from the UCSC Genome Browser [33]. Windows were labeled as containing a respective repeat type if there was any overlap with the corresponding annotation from RepeatMasker.

### 2.8 Evolutionary analyses

To construct a cladogram showing the evolutionary relationships between species, we use the phyloT v2 visualization tool. This program generates phylogenetic trees based on the NCBI taxonomy or Genome Taxonomy Database using a list of taxonomic names, identifiers, or protein accessions [34].

## 3 Results

### 3.1 Approach

Our approach, MORALE, is a straightforward generalization of the Deep CORAL [15] method, applied to TF binding data. In the following we briefly describe the method:

We assume labeled (sequence) data from *n* different source domains, and unlabeled data from a target domain. In our application, the domains are species. We also have an encoder model (see **Section 2.2**), mapping input sequences *x*^*i*|*j*^ to vector embeddings *z*^*i*|*j*^ ∈ ℝ^*d*^, where *i* = 1, …, *m*_*j*_ and *j* = 1, …, *n* + 1 and *m*_*j*_ is the overall number of sequences in the *j*-th domain. The domain index *j* runs to *n* + 1 because the target domain is included. Vector representations, in turn, are input to a classification model predicting the sequence label (e.g., whether the input sequence binds a specific TF) with an associated loss (e.g., cross entropy) that we denote by ℒ_label-classification_.

Given a number of sequence representations 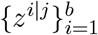 for a *b < m*_*j*_ we denote the sample estimate of the *z*’s mean by 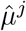 and of the covariance matrix by 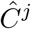. For a mini-batch containing multiple examples/sequences of each domain/species, the MORALE loss is then calculated as

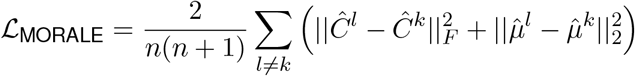

quantifying the difference in first and second moments between all domain pairs. Here || *·* ||_*F*_ denotes the Frobenius (Euclidean / *L*^2^) norm for matrices. In our analyses we estimate the moments 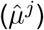 and 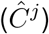 for the (ℒ_[MORALE]_) loss by utilizing the ‘background’ portion of the mini-batch (described in **Section 2.1**), which contains an equal number of unlabeled sequences sampled from each domain (source(s) and target). For source domains, the vector representations 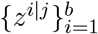 are used for label classification.

The MORALE loss is then added to the label classification loss, encouraging a vector representation of examples/sequences that is moment-aligned across domains:

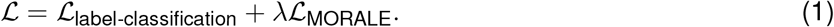

We note gradient reversal layers (GRL) are a successful adversarial learning approach to encourage domain-invariant representations [35–37]. Briefly, a domain classifier and associated classification loss for predicting embeddings’ domains (e.g., cross entropy) is added to the model for an overall loss of

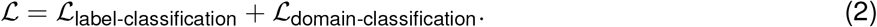

Backpropagated gradients from the corresponding domain classification loss ℒ_domain-classification_ are penalized by a factor of *™λ* (for *λ >* 0, therefore “gradient reversal”), encouraging vector representations that are not informative about the domain of origin.

In our analyses we compare MORALE (**Equation 1**) with the GRL approach (**Equation 2**) and find it generally outperforms GRL. We also note that MORALE does not require additional parameters on top of the encoder and label classification model, whereas GRL requires additional design and parameterization of the domain classifier. **Figure 1** summarizes our approach, also in the context of gradient reversal and naive classification without domain adaptation.

**Figure 1:**
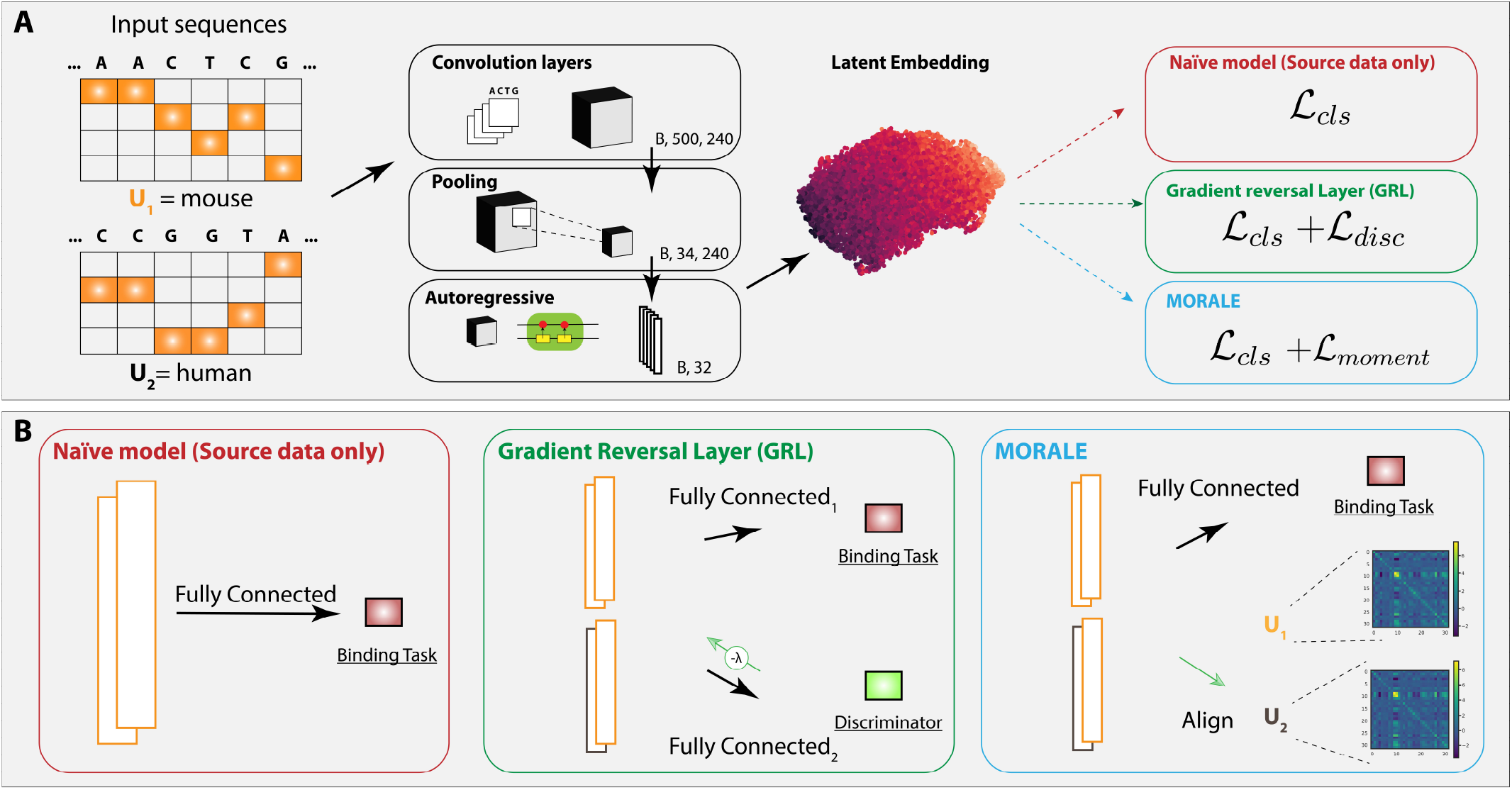
Schematic overview of domain adaptation framework with MORALE compared to leading alternatives. **(A)** We one hot encode input sequences from the source and target domains (i.e., human and mouse) such that we can create a latent embedding after a set of convolution layers, pooling, and autoregressive components. The embedding is then used for all pertinent tasks downstream (classification and domain-adaptive). We evaluate on three procedures: (1) Training a naive model on source data only, to predict in the target domain, (2) Leveraging an adversarial, discriminator approach to penalize learned gradients, and (3) Leveraging a loss that operates on the moment alignment of all domains of data, which is added to the total loss. In **(B)** we describe the setup of the different approaches in more detail, including that of MORALE: (1) Under the naive, source-only model, we input the relevant part of the batch (the embedded features coming from the source domain) into a classifier head to predict the label for the corresponding sites. After training, this can be evaluated on the target domain. (2) The GRL approach adds a separate branch to the overall scheme. We still predict labels during training on the source data, but now also feed in the source and target data (in the batch) to a classifier whose purpose is to predict the domain (*U*) the data comes from. This learned gradient is then penalized to encourage the learning of an invariant representation for downstream target evaluation. (3) Finally, we showcase MORALE. We use one branch to predict labels on the source features during training. We also align the moments of the data between the source and target in an intermediary stage. In order to have an effect on training, this moment alignment loss is added to the overall model loss during training.

### 3.2 Cross-species TF-binding prediction between human and mouse

#### 3.2.1 MORALE improves cross-species TF binding prediction performance

First, we applied our framework to re-analyze a dataset introduced by Cochran et al. [13]. In this scenario, the binding of four TFs (CTCF, HNF4*α*, RXRA, and CEBPA) was assayed in liver tissue samples from humans and mice. This makes for two species/domains, and we assess TF binding site prediction for each species as target, and the other as source. See the Methods section for details. Test set performances are summarized in **Figure 2**. For each TF, we report the performance of the source-trained model on the target data labeled as “source” (i.e., no domain adaptation, no access to the target domain); the target-trained model on the target data labeled as “target” (i.e., learned within the target domain and with access to binding labels on that domain); and two approaches for domain adaptation (with access to vector embeddings on the target domain, but without access to binding labels on the target domain): our method (labeled “MORALE”) and domain classification task with gradient reversal layer (labeled “GRL”).

**Figure 2:**
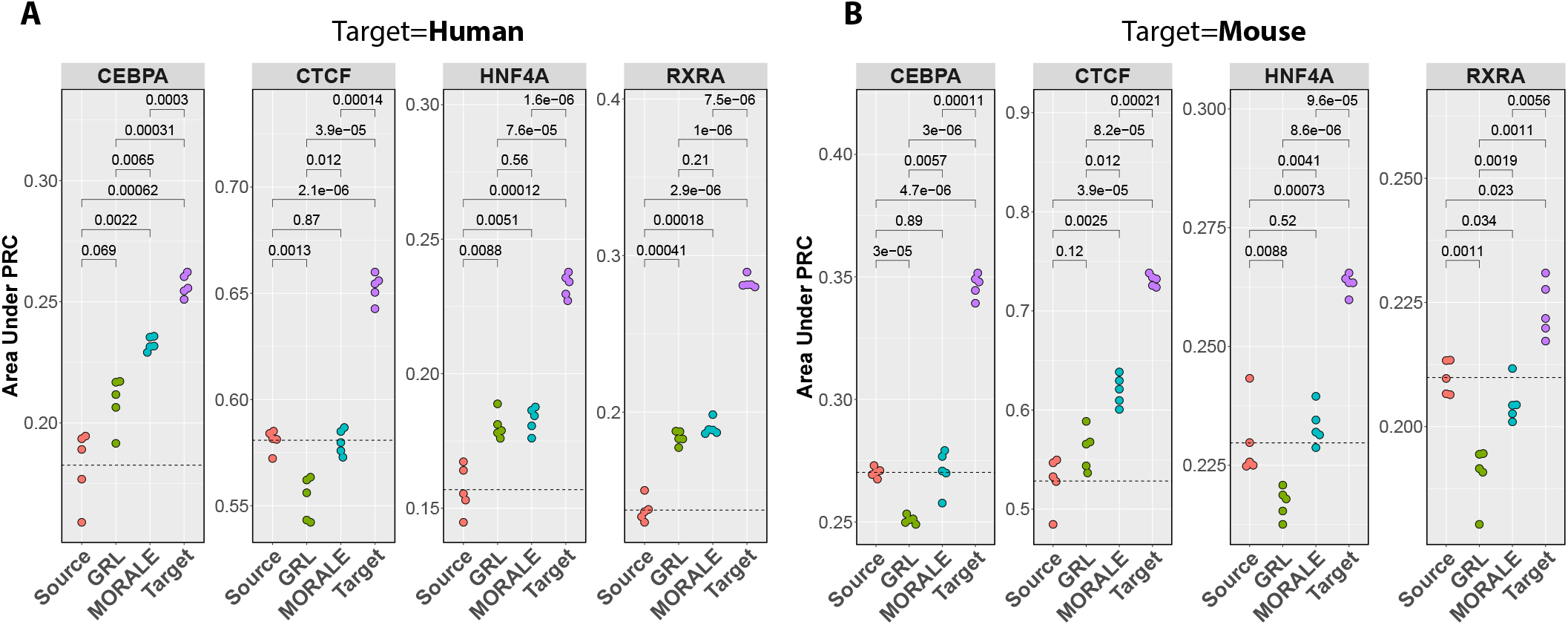
Moment alignment improves cross-species TF binding site predictions. For four TFs and human as the target species, prediction performance is shown for four models: (1) Source-trained on target (red), (2) gradient reversal (green), (3) moment alignment (blue), and (4) and target-trained on target (purple). **(A)** We display the results across the four transcription factors when adapting mouse model to human data. MORALE outperforms or matches the GRL performance in each case, while not suffering from degradation. **(B)** The same as in (A), but in the other adaption direction, human adapted models to mouse. We find that the degradation is persistent in this study under the scope of using gradient reversal. MORALE is able to at least meet source-trained baselines, or outperform the GRL.

As expected, we overall find that no domain adaptation (i.e., “source”) performs the worst, training on target (i.e., “target”) performs the best, and domain adaptations are in-between. Interestingly, we find that using human as target species for CTCF, the GRL approach performs worse than no domain adaptation, while MORALE does not suffer from this drop in performance. Likewise, using mouse as target, GRL performs worse than no domain adaptation in three (CEBPA, HNF4*α*, RXRA) of the four TFs; again, MORALE does not experience a corresponding performance drop. In addition, for the TFs where GRL domain adaptation outperforms no domain adaptation, MORALE either matches or exceeds GRL’s performance. The exact numerical values for the comparisons in can be found in **Table 1** under the column “auPRC”, we exclude the target model as none of the source or source-adapted models come close to its performance.

**Table 1:** We display the true positives (TPs), false positives (FPs), false negatives (FNs) with the auPRC across four TFs and three methods in this study. In order to generate this confusion matrix across five-folds of data, we average the sigmoid value over the folds and compute the relevant prediction types w.r.t. the ground truth label, percentages are with respect to the complete dataset. We observe that the GRL more often predicts positives than either the source model, or MORALE, however this comes at much higher occurrence of FPs, leading to performance degradation as in CTCF when the target is human. MORALE, on the other hand, takes a much more conservative step towards the target model, finding a middle ground that strictly results in improvement or meeting baselines.

| Target=Human |  |  |  |  |  |  |  |  |  |  |  |  |
| --- | --- | --- | --- | --- | --- | --- | --- | --- | --- | --- | --- | --- |
| TF | TPs (%) |  |  | FPs (%) |  |  | FNs (%) |  |  | auPRC |  |  |
|  | GRL | MORALE | Source | GRL | MORALE | Source | GRL | MORALE | Source | GRL | MORALE | Source |
| CTCF | **0.413** | 0.402 | 0.401 | 1.847 | **1.384** | 1.397 | **0.068** | 0.079 | 0.08 | 0.553 | 0.58 | **0.581** |
| CEBPA | 0.542 | 0.545 | **0.547** | 10.436 | **9.77** | 11.938 | 0.048 | 0.045 | **0.044** | 0.209 | **0.233** | 0.183 |
| HNF4A | **0.605** | 0.578 | 0.574 | 9.137 | **8.496** | 10.806 | **0.136** | 0.164 | 0.168 | 0.181 | **0.183** | 0.157 |
| RXRA | **1.133** | 1.035 | 1.076 | 9.981 | **8.023** | 13.181 | **0.468** | 0.566 | 0.526 | 0.184 | **0.19** | 0.137 |
| Average | **0.673** | 0.64 | 0.649 | 7.85 | **6.918** | 9.33 | **0.18** | 0.213 | 0.205 | 0.282 | **0.296** | 0.264 |

| Target=Mouse |  |  |  |  |  |  |  |  |  |  |  |  |
| --- | --- | --- | --- | --- | --- | --- | --- | --- | --- | --- | --- | --- |
| TF | TPs (%) |  |  | FPs (%) |  |  | FNs (%) |  |  | auPRC |  |  |
|  | GRL | MORALE | Source | GRL | MORALE | Source | GRL | MORALE | Source | GRL | MORALE | Source |
| CTCF | **0.7** | 0.699 | 0.697 | 3.937 | **3.237** | 3.591 | **0.042** | 0.044 | 0.046 | 0.56 | **0.62** | 0.528 |
| CEBPA | 0.988 | **0.996** | 0.952 | 7.158 | 6.66 | **5.732** | 0.335 | **0.328** | 0.372 | 0.251 | **0.271** | 0.27 |
| HNF4A | 0.824 | **0.828** | 0.813 | 11.026 | 10.388 | **9.802** | 0.122 | **0.118** | 0.133 | 0.217 | **0.233** | 0.23 |
| RXRA | 0.831 | 0.827 | **0.843** | 17.891 | **16.039** | 19.208 | 0.087 | 0.091 | **0.075** | 0.19 | 0.205 | **0.21** |
| Average | 0.836 | **0.837** | 0.826 | 10.003 | **9.081** | 9.583 | 0.146 | **0.145** | 0.156 | 0.304 | **0.332** | 0.31 |

Upon laying out the confusion matrix (**Table 1**) to understand what types of predictions are being made to contribute to the model performance, we observe that in the mouse-to-human direction the GRL model often makes more positive predictions overall, but also has disproportionately more false positive predictions compared with MORALE. This leads to comparatively worse performance, especially for CTCF. MORALE, on the other hand, has lower false positive rates (e.g., 1.38% versus the GRL’s 1.85% for CTCF) while suffering from only slightly increased false negative rates (0.079% vs. 0.068%, again for CTCF). In the human-to-mouse direction this is also the case for CTCF. For CEBPA and HNF4*α* MORALE has more true positives and less false positives than GRL, and consequently shows better performance. For RXRA MORALE’s performance is close to the source model, while GRL performs somewhat worse.

From these analyses, we conclude that MORALE improves over GRL for cross-species TF binding prediction on this dataset. Next, we compare models and the quality of their predictions in more detail.

#### 3.2.2 MORALE improves cross-species TF binding prediction quality

To quantify/inspect the quality of TF binding predictions and compare MORALE and GRL domain adaptation approaches, we take the following approach. Taking the target-trained model as a reference, we compare how well MORALE and GRL “adapt” the source-trained model to the target-trained model. We use two metrics to quantify this concept, both of which rely on importance scores generated by post-hoc attribution analysis (see Methods section for details, **Section 2.6**). Briefly, for a given model, these scores summarize the importance of a sequence position. We use those scores to quantify the quality of the GRL/MORALE models in two ways.

First, we compare the correlation of GRL/MORALE importance scores with importance scores of the “target” model. Specifically, we focus on sequences where the “source” model disagrees with the target model: on differential false positives (dFPs), where the “source” model wrongly predicts TF binding but the target model does not; and on differential false negatives (dFNs), where the “source” model wrongly predicts no TF binding, but the “target” model correctly predicts a binding event. **Figure 3** shows contour plots of correlation coefficients between GRL/MORALE scores and “target” scores, stratified by TF and by type of sequence (dFP/dFN). Higher correlation means that the corresponding domain adaptation method better reproduces the importance of the target-trained model and is desirable.

**Figure 3:**
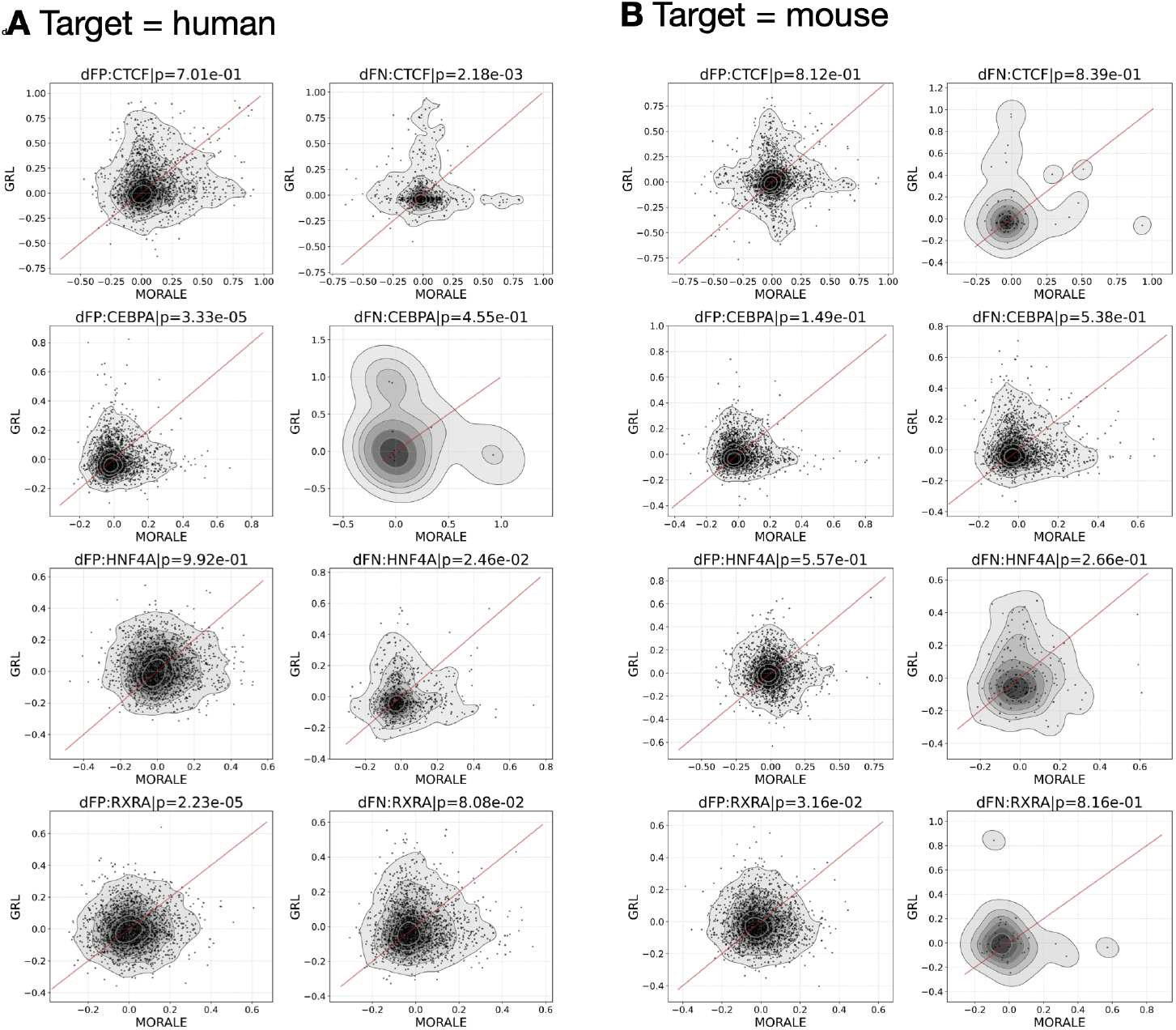
Pearson correlation coefficients between MORALE’s importance scores and the “target” model (x-axes) and GRL’s importance scores and the “target” model (y-axes) for differentially false positive sites (dFPs) and differentially false negative sites (dFNs), see text for details. Panel A shows the analyses for mouse-to-human and panel B for human-to-mouse. P-values for a one-sided Wilcoxon Rank Sum Test are indicated for each plot.

We see that for the mouse-to-human direction and for dFP sequences, MORALE importance scores are significantly more correlated with the target model than GRL scores for CEBPA and RXRA, whilst things look even for the CTCF and HNF4*α* comparisons; for dFN sequences, MORALE’s correlation is significantly closer to the target models for RXRA and HNF4*α*. For human-to-mouse and dFPs, MORALE importance scores have higher correlation with target scores for RXRA, while the other comparisons (also for dFN sequences) are not significant. Overall, this analysis demonstrates that MORALE can improve upon GRL in terms of highlighting meaningful sequence positions after domain adaptation, while we have not observed the reverse (GRL improving over MORALE).

Second, based on calculated attribution scores via expected integrated gradients, we quantified sequence motifs highlighted by the different models for TF binding prediction using TF-MoDISCo ([31]) (see Methods section for details, **Section 2.6**). **Figure 4** summarizes our findings. In panel A we see that for CTCF binding sites the most-frequently found motif of the target model corresponds to CTCF, which is recapitulated in the MORALE domain adaptation model. GRL domain adaptation, like the source-only trained model, finds a larger fraction of motifs corresponding to other TFs. In panels B to E we see that the target-trained, MORALE, and the source-only trained model find CTCF as their most-frequent motif matches for CTCF-bound sites. Surprisingly, GRL domain-adapts the source-only trained model in a way that motifs that poorly match CTCF become more frequent.

**Figure 4:**
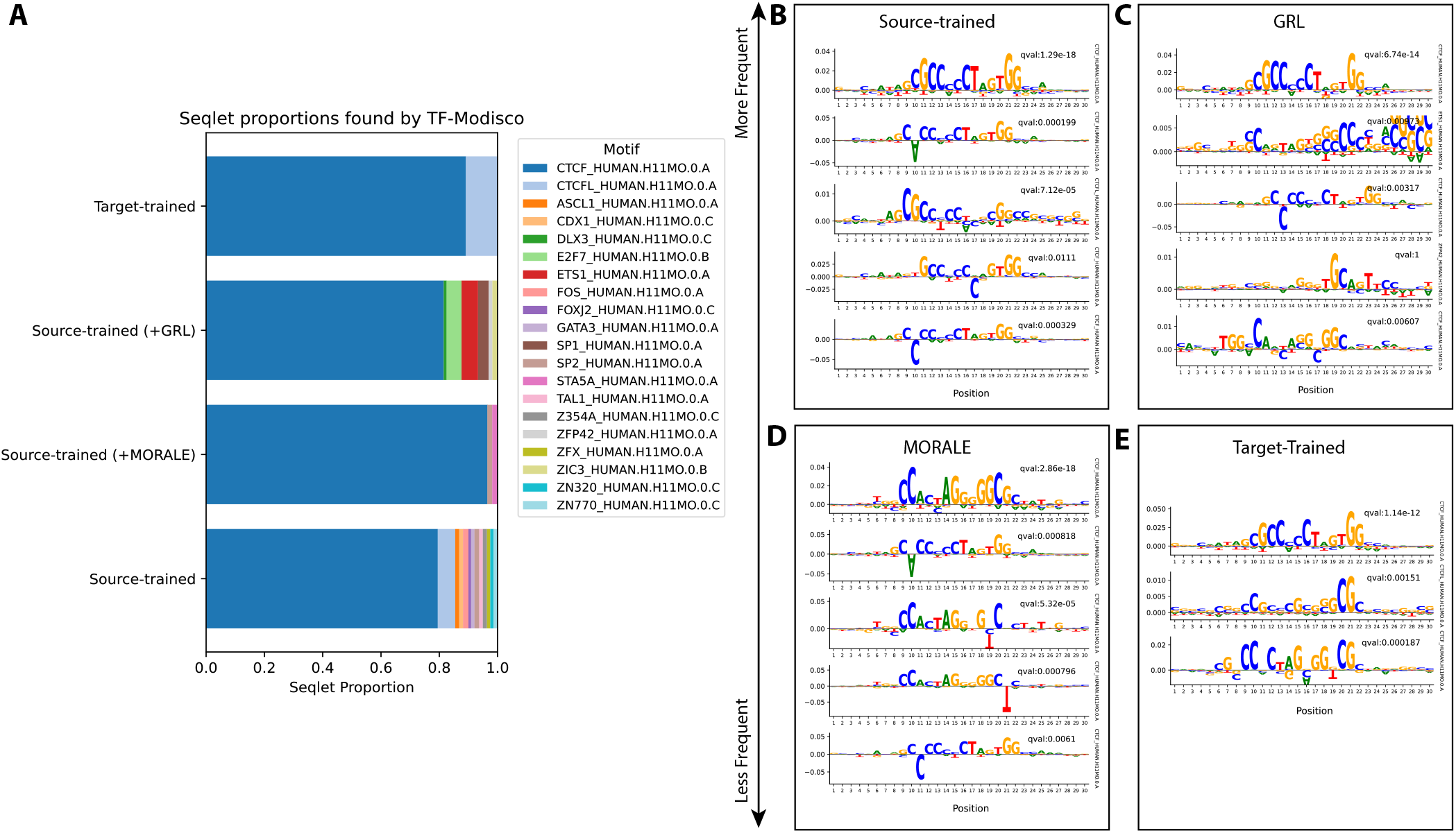
MORALE discovers de-novo motifs more similar to CTCF. Using calculated attribution scores across source, source-adapted, and target models, we compare the de-novo motifs found across 2000 randomly samples bound sites for CTCF. We display the output of TF-MoDISCo in the following format. In **(A)** we show the proportions of all found motifs across the four models, notably, the target model (human-on-human) only finds CTCF (and CTCFL, a paralog). The source-trained model, along with all source-adapted models, in the majority, do find CTCF primarily, however both find other motifs in these CTCF bound sites at a higher proportion than MORALE, which nearly only report CTCF. In **(B)** We display the top 5 de-novo motifs found by the source trained model, with annotated p-values and the corresponding TomTom matches on the right, y-axes. The same is done for **(C), (D)**, and **(E)**. The top match of MORALE strongly resembles the established CTCF motif with a significant q-value.

Finally, we note that Cochran et al. [13] have reported that the GRL domain-adapted model reduces over-prediction of TF binding sites in the mouse-to-human direction. Therefore, we explored the performance of GRL, MORALE and the source-only trained base model (“source”) on test data overlapping LINE and SINE elements. Results are summarized in **Table** 2. Indeed, we observe that the GRL domain-adapted model outperforms both MORALE and “source” for HNF4*α* and especially for RXRA in this direction. Nevertheless, for CTCF and CEBPA MORALE does outperform GRL. Further on, in the human-to-mouse direction, MORALE outperforms GRL for all four TFs and has the best average performance for three of the four.

Overall, comparing MORALE with GRL on this dataset we conclude that MORALE outperforms GRL for some cases, while it rarely performs worse. This renders MORALE a robust domain adaptation method in this setting.

### 3.3 Learning TF-binding in human by leveraging data across five mammals

Next, we explored whether multiple species, together with domain adaptation, can improve TF binding prediction in human. Using ChIP-seq data from rhesus macaque, rat, mouse, and dog liver, we evaluate generalization to human target/test data, comparing three approaches: training solely on human data (the best case, target-on-target model, but trained on a single domain), training on multi-species data without adaptation, and training on multi-species data with MORALE’s correction for moments. We study four TFs, FOXA1, HNF4*α*, HNF6, and CEBPA. Results are summarized in **Figure 5**. We observe two notable behaviors under the scope of working with five mammalian domains for TF binding prediction. The first, as in **Figure 2**, the source-adapted models, leveraging information from two domain, cannot approach the performance of the target-on-target model in any case. However, now when training a plain model under the scope of five mammals, we are able to outperform the target-on-target model on all TFs under the scope of the human species being the target species. The next observation is with regards to MORALE, as when applied to the joint model, we are still able to increase performance past the joint model. MORALE is the top performer for each TF under this setting, indicating its broad application to encourage a learned, invariant representation in the case of multiple (*>* 2) domains.

**Figure 5:**
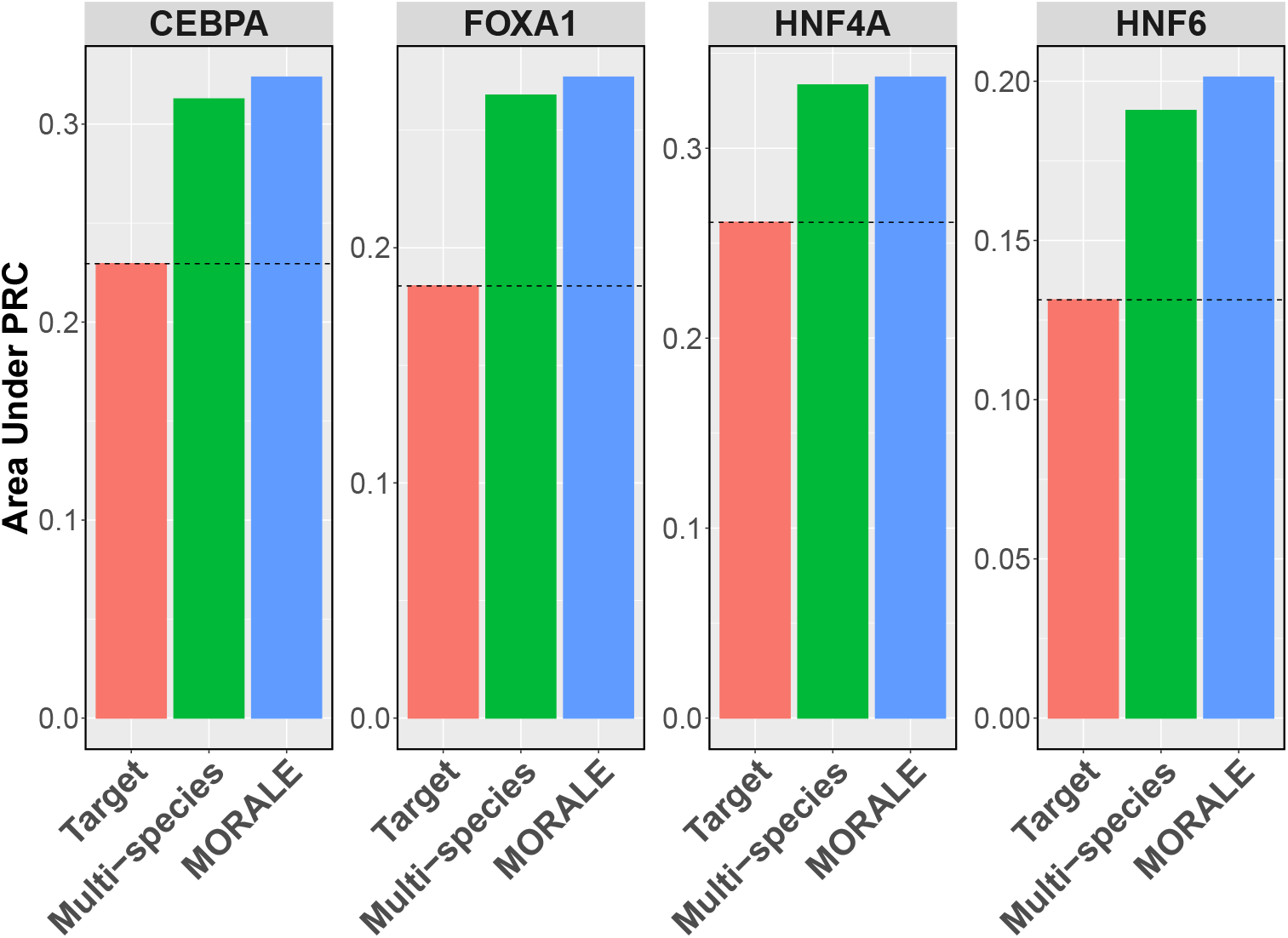
MORALE attains higher performance when training on multiple source species and predicting in human as the target. Across the four transcription factors tested, leveraging MORALE unilaterally increases performance — compared to no domain adaptation at all. Of note, unlike the two-species case, training on multiple source species allows us to outperform the human-on-human model, previously acting as an upper bound not attainable.

**Table 2:** We display the performance (auPRC) for windows that overlap both SINE and LINE repeats, and their average. We observe that MORALE performs competitively in the performance across windows that contain species-specific repeats in the mouse-to-human direction, and outperforms the GRL in the human-to-mouse direction. Notably, for HNF-4*α* and RXR*α*, the GRL does a convincly better job windows that overlap SINE elements in human, which would be Alus in the majority.

| Target=Human |  |  |  |  |  |  |  |  |  |
| --- | --- | --- | --- | --- | --- | --- | --- | --- | --- |
| TF | LINE |  |  | SINE |  |  | Average |  |  |
|  | GRL | MORALE | Source | GRL | MORALE | Source | GRL | MORALE | Source |
| CTCF | 0.149 | **0.173** | 0.17 | 0.181 | **0.191** | 0.177 | 0.165 | **0.182** | 0.173 |
| CEBPA | 0.039 | **0.042** | 0.034 | 0.04 | **0.04** | 0.027 | 0.039 | **0.041** | 0.031 |
| HNF4A | **0.043** | 0.04 | 0.03 | **0.048** | 0.034 | 0.023 | **0.046** | 0.037 | 0.026 |
| RXRA | **0.058** | 0.056 | 0.036 | **0.062** | 0.052 | 0.029 | **0.06** | 0.054 | 0.033 |
| Average | 0.072 | **0.078** | 0.068 | **0.083** | 0.079 | 0.064 | 0.077 | **0.079** | 0.066 |

| Target=Mouse |  |  |  |  |  |  |  |  |  |
| --- | --- | --- | --- | --- | --- | --- | --- | --- | --- |
| TF | LINE |  |  | SINE |  |  | Average |  |  |
|  | GRL | MORALE | Source | GRL | MORALE | Source | GRL | MORALE | Source |
| CTCF | 0.103 | **0.124** | 0.111 | 0.102 | **0.129** | 0.104 | 0.102 | **0.126** | 0.107 |
| CEBPA | 0.077 | 0.08 | **0.082** | 0.097 | 0.103 | **0.108** | 0.087 | 0.092 | **0.095** |
| HNF4A | 0.04 | 0.043 | **0.045** | 0.061 | **0.063** | 0.062 | 0.051 | **0.053** | 0.053 |
| RXRA | 0.027 | **0.029** | 0.026 | 0.041 | **0.042** | 0.035 | 0.034 | **0.036** | 0.03 |
| Average | 0.062 | **0.069** | 0.066 | 0.075 | **0.084** | 0.077 | 0.068 | **0.077** | 0.071 |

Given this result, we move towards a working understanding of the effect of the domain contribution based on the individual species. We lay out the phylogenic tree of evolutionary distance at the top of **Figure 6**, constructed with phyloT (see **Section 2.8** for details). We perform a species-holdout analysis when the target is human and showcase the auPRC results in panel A. Given the tree, we would expect, in order, rhesus macaque, mm10, rn7, and finally canFam6 to have a series of decreasing effects on performance. Nevertheless, we observe a slight deviation from the expectation in these findings. Across three of the four TFs (CEBPA, FOXA1, and HNF4*α*), as expected, the rhesus macaque holdout has the largest impact on predicting on human. The other holdouts more or less have an impact, but may or may not follow such a trend in the expected cost of holding each out; we observe for HNF6 especially, the expected evolutionary trend does not hold. Interestingly, for HNF6, the rn7 holdout tends to have just as large of an impact on the performance when the target is human (rn7 tends to be enriched in bound sites when compared to other species for the same TF, which can be seen in **STab. 8)**.

**Figure 6:**
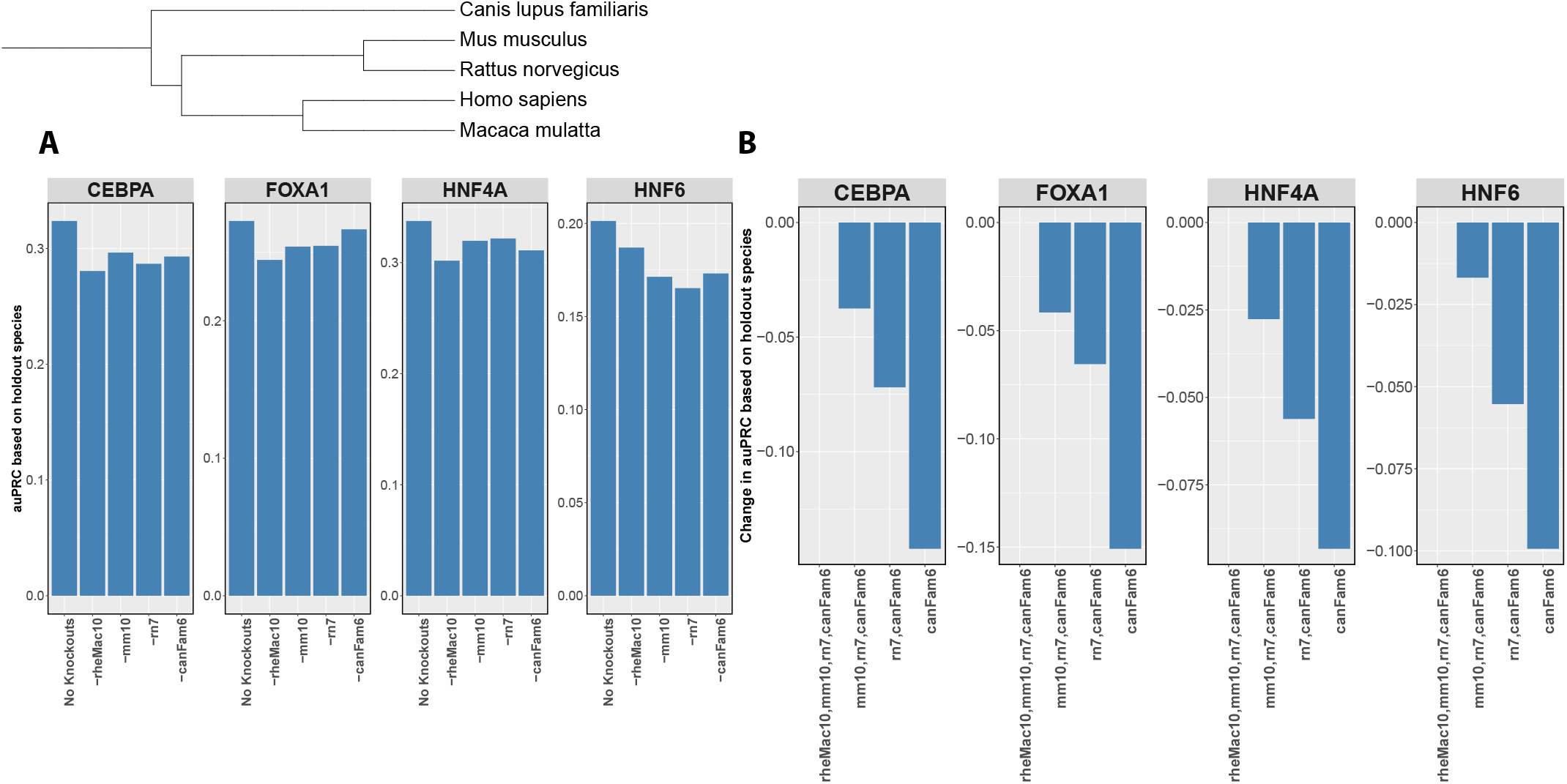
Species contribution to the overall boost in mutli-species results shows to be differential in nature, still ultimately resulting in benefit. We seek to quantify the effect of holding out species: (1) individually, and (2) in groups, to understand how it changes model performance under the scope of using MORALE for domain-adaption. In **(A)** we present the results for the per-species holdout performance. ‘No knockout’ describes the model performance when all source species are used, with human as the target species. From left to right, we proceed to holdout a single species from the overall source species to determine its holdout’s affect on performance. In **(B)** we display the results for holding our groups of species at a time. From left to right, we display a monotonically decreasing set of source species from all included (i.e., leftmost) to just a singular species (i.e., rightmost). This showcases that the number of species included in training set does aide in overall model performance.

We move to a group holdout analysis in panel B. We perform successive group holdouts to showcase that the effect of training of multiple species aids in performance over the target-on-target model. From left-to-right we holdout: (1) No species, (2) rhesus macaque, (3) rhesus macaque, and mouse, (4) rhesus macaque, mouse, and rat. We see parallel observations in the group holdout setting that we observed in panel A, each successive holdout contributing to the drop in performance — emphasizing the importance that the multiple species aide to the overall performance over the target-on-target model.

## 4 Discussion

In this study, we develop and apply a new framework, MORALE, for domain adaptation in the context of generalizing TF binding across species. The MORALE approach encourages a domain-invariant sequence representation, and we show it improves upon an adversarial approach in a two-species case, and that it can successfully leverage additional information in a multi-species case. Specifically, we show that MORALE is able to avoid performance degradation observed by an adversarial approach in both adaptation directions, that is, when predicting TF binding in mouse liver, based on human data, and vice versa. In addition, we show that using TF binding data in five species noticeably improved TF binding site prediction in human liver. In contrast to current approaches like gradient reversal layers, MORALE is not adversarial and does not require an adversarial component in the network model. Instead, MORALE uses moment alignment [15] and aligns the learned moments of sequence embeddings for data across all pairs of domains. MORALE integrates easily with other approaches that utilize embedding layers and does not require extra parameters or decisions in model design, like creating an adversarial classifier. We believe this is a meaningful improvement over the status quo in TF binding prediction, because identifying precise, evolutionarily conserved sequence patterns that enhance and complement widely used TF binding site motifs can advance our understanding of the gene regulatory code. Specifically, it is attractive to address TF binding from a causal modeling perspective. In this paradigm, domain adaptation techniques like MORALE are needed to construct identifiable latent sequence embeddings that typically underlie this type of analysis.

## 5 Conclusion

MORALE is a domain adaptation method for cross-species sequence-based modeling using representation learning in computational genomics. It allows for flexibility in addressing species-specific differences, and it is geared toward stable model generalization. MORALE helps reveal learned invariant sequence representations in regulatory genomics and advances the study of TF-DNA binding.

## 6 Acknowledgments

We thank Shaun Mahony for the helpful discussions and the processed data used in this work.

## 9 Data availability

See the methods section. All of the data used in this analyses can be found publicly; the accession IDs are listed in **STabs**. 7, and 8.

## 10 Supplement

**Figure 7:**
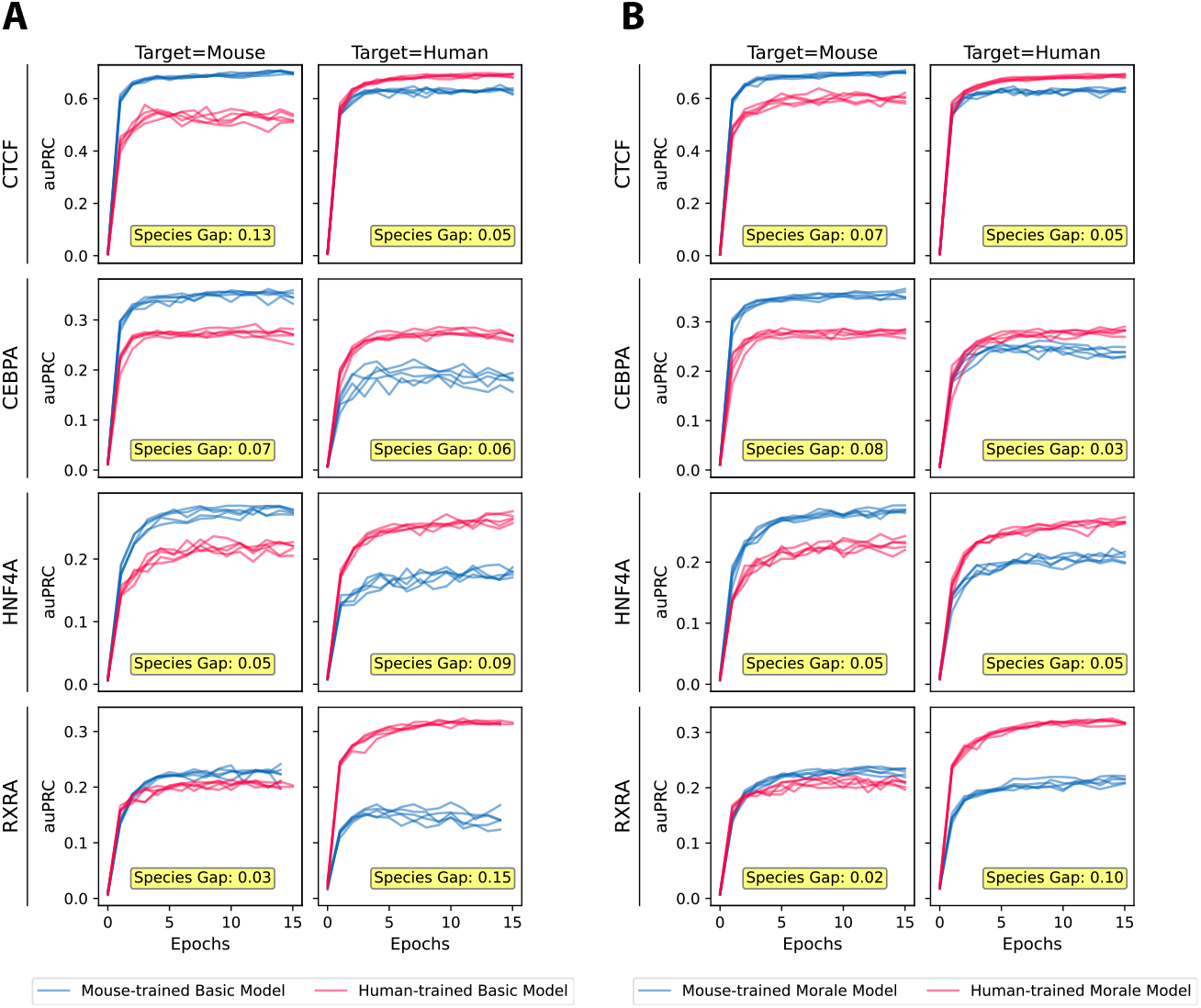
The species gap is closed during training when using MORALE. We display the training, in the two-species case, of the five-fold cross validation performance over 15 epochs between the basic model, and MORALE. The gap between the peak performance between the source-on-target and the target-on-target models are annotated at the bottom of each plot. In **(A)** we have the basic model across each TF-target pair, and in **(B)** we display the same information, but with MORALE.

**Table 3:**
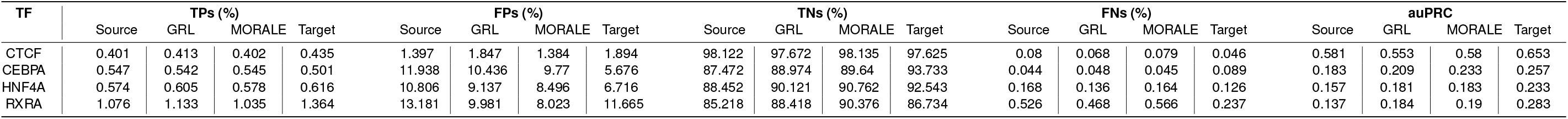
The confusion matrix for mouse-adapted models when evaluating of the test set (Chr2) from human. The table displays the percentage of true positives (TPs), false positives (FPs), true negatives (TNs), and false negatives (FNs) for each TF. We include them as a ratio over all the windows in the test set and attach the auPRC value based on model type.

**Table 4:**
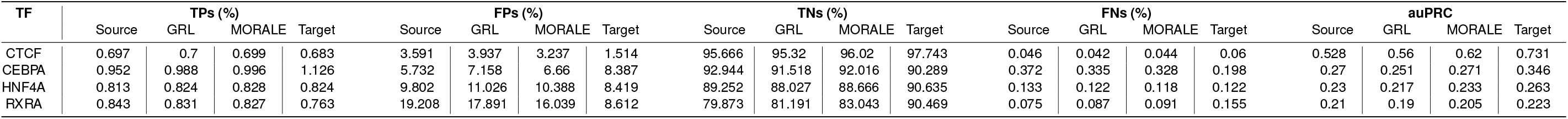
The confusion matrix for human-adapted models when evaluated on the test set (Chr2) from mouse. The table displays the percentage of true positives (TPs), false positives (FPs), true negatives (TNs), and false negatives (FNs) for each TF. We include them as a ratio over all the windows in the test set and attach the auPRC value based on model type.

**Table 5:**
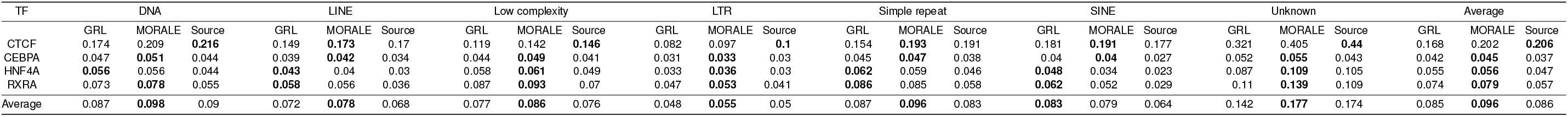
We compare across all repeat types (with at least 500 instances in our test (human) chromosome, Chr2) in the source (mouse) genome between the two mouse-adapted models. The last row is the average auPRC across all repeat types

**Table 6:**
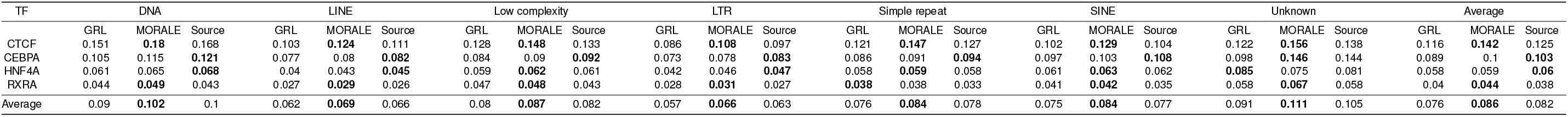
We compare across all repeat types (with at least 500 instances in our test (mouse) chromosome, Chr2) in the source (human) genome between the two human-adapted models. The last row is the average auPRC across all repeat types

**Table 7:** The binding site information for the four transcription factors used in the two-species case. The following quantities are listed: the number of peaks called across the entire genome; the number of called peaks within the filtered window set, merged if within 500 bp of each other; the number of windows in the filtered window set labeled bound due to peak overlap; the fraction of the filtered window set labeled bound; and the database accession ID (ENCODE, GEO, or ArrayExpress). The size of the filtered window sets for the mouse and human genomes were 41883806 and 48742577, respectively.

| TF | Species | Raw Peaks | Filtered Peaks | Bound Windows | Frac. Bound | Accession ID |
| --- | --- | --- | --- | --- | --- | --- |
| CTCF | Mouse | 32006 | 28943 | 296117 | 0.71% | ENCSR000CBU |
|  | Human | 29067 | 26477 | 270100 | 0.55% | ENCSR911GFJ |
| CEBPA | Mouse | 62636 | 48812 | 566945 | 1.35% | E-TABM-722 |
|  | Human | 32243 | 28545 | 298066 | 0.61% | E-TABM-722 |
| HNF4A | Mouse | 44800 | 36540 | 415846 | 0.99% | E-TABM-722 |
|  | Human | 42766 | 34714 | 387077 | 0.79% | E-TABM-722 |
| RXRA | Mouse | 46443 | 33751 | 404284 | 0.97% | GSM1299600 |
|  | Human | 95085 | 71032 | 854289 | 1.75% | ENCSR098XMN |

**Table 8:**
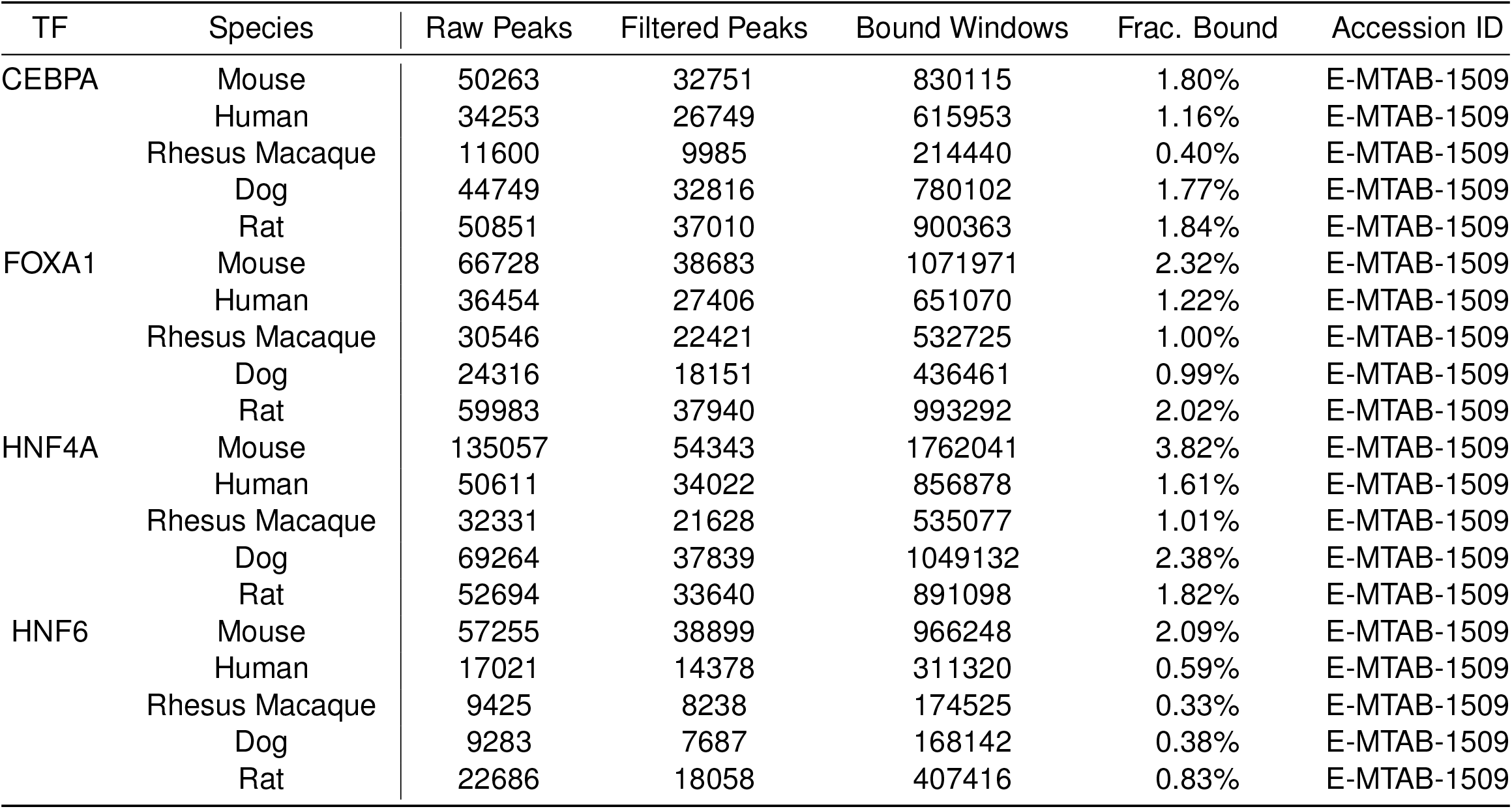
The binding site information for the four transcription factors used in the multi-species case. The following quantities are listed: the number of peaks called across the entire genome; the number of called peaks within the filtered window set, merged if within 1000 bp of each other; the number of windows in the filtered window set labeled bound due to peak overlap; the fraction of the filtered window set labeled bound; and the database accession ID (ArrayExpress).

**Figure 8:**
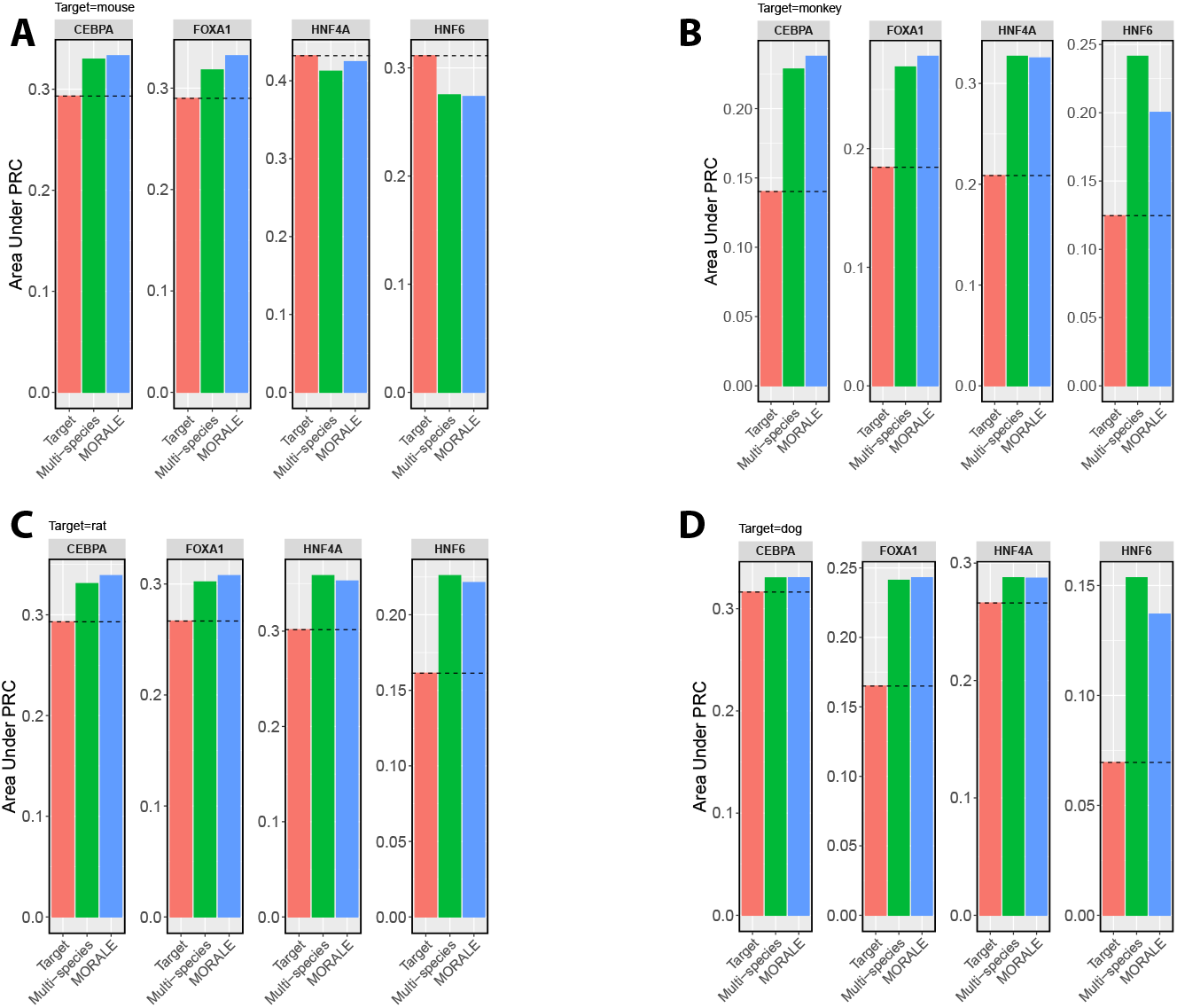
We display performance across all targets in the multi-species case. In **(A)** we show the performance across all 4 transcription factors when the target is mouse, **(B)** displays monkey, **(C)** displays rat, and **(D)** display dog.

**Figure 9:**
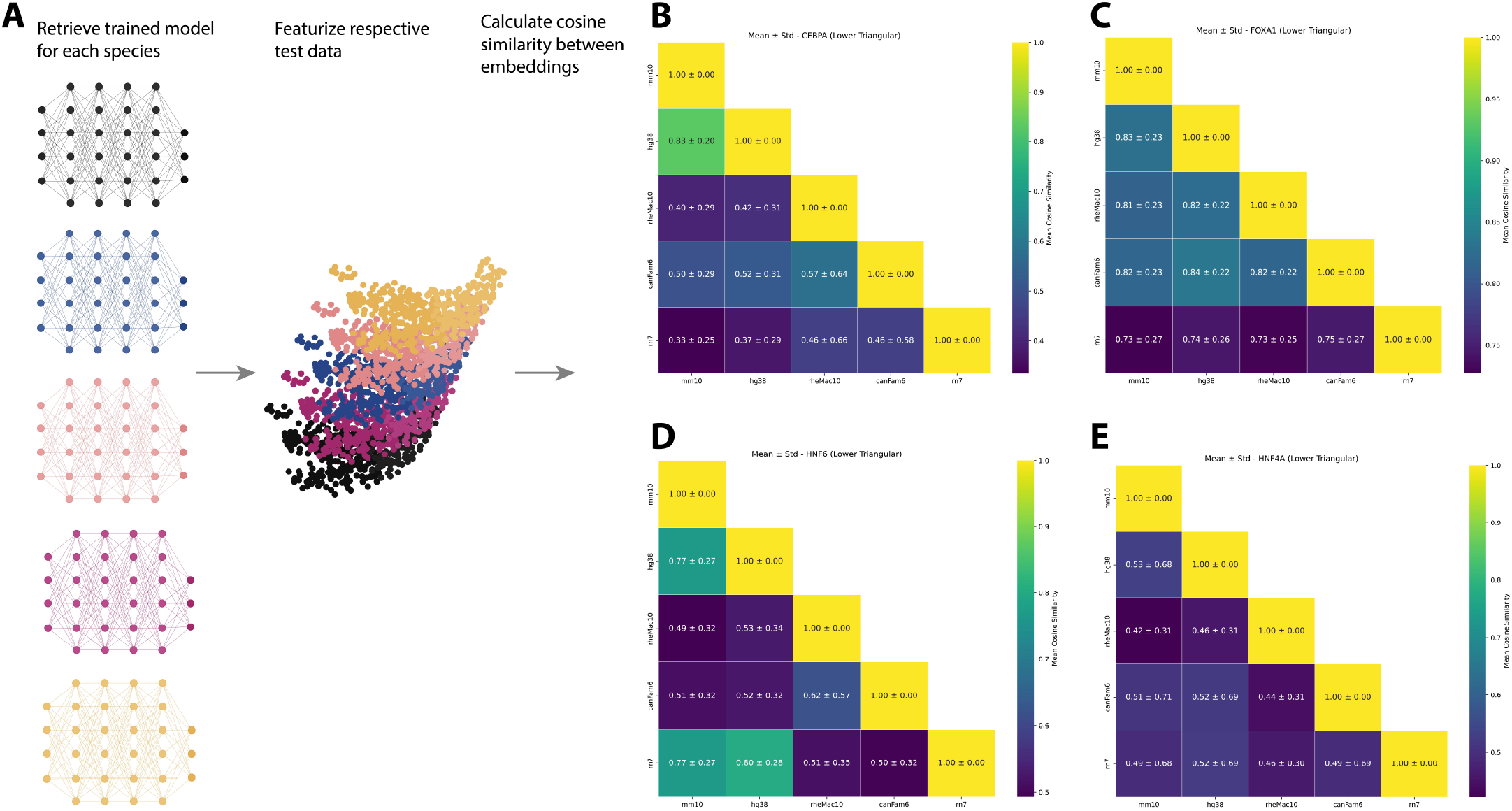
We construct heatmaps based to understand relatedness of learned embeddings through the different species models. We do so for each TF under study in the multi-species case. For each TF we use the models trained to predict in each species and run the test data through the feature extractor in our models to capture the embeddings. In (**B**) we should the lower triangular for CEBPA, (**C**) FOXA1, (**D**) HNF6, and (**E**) HNF4*α*.

**Figure 10:**
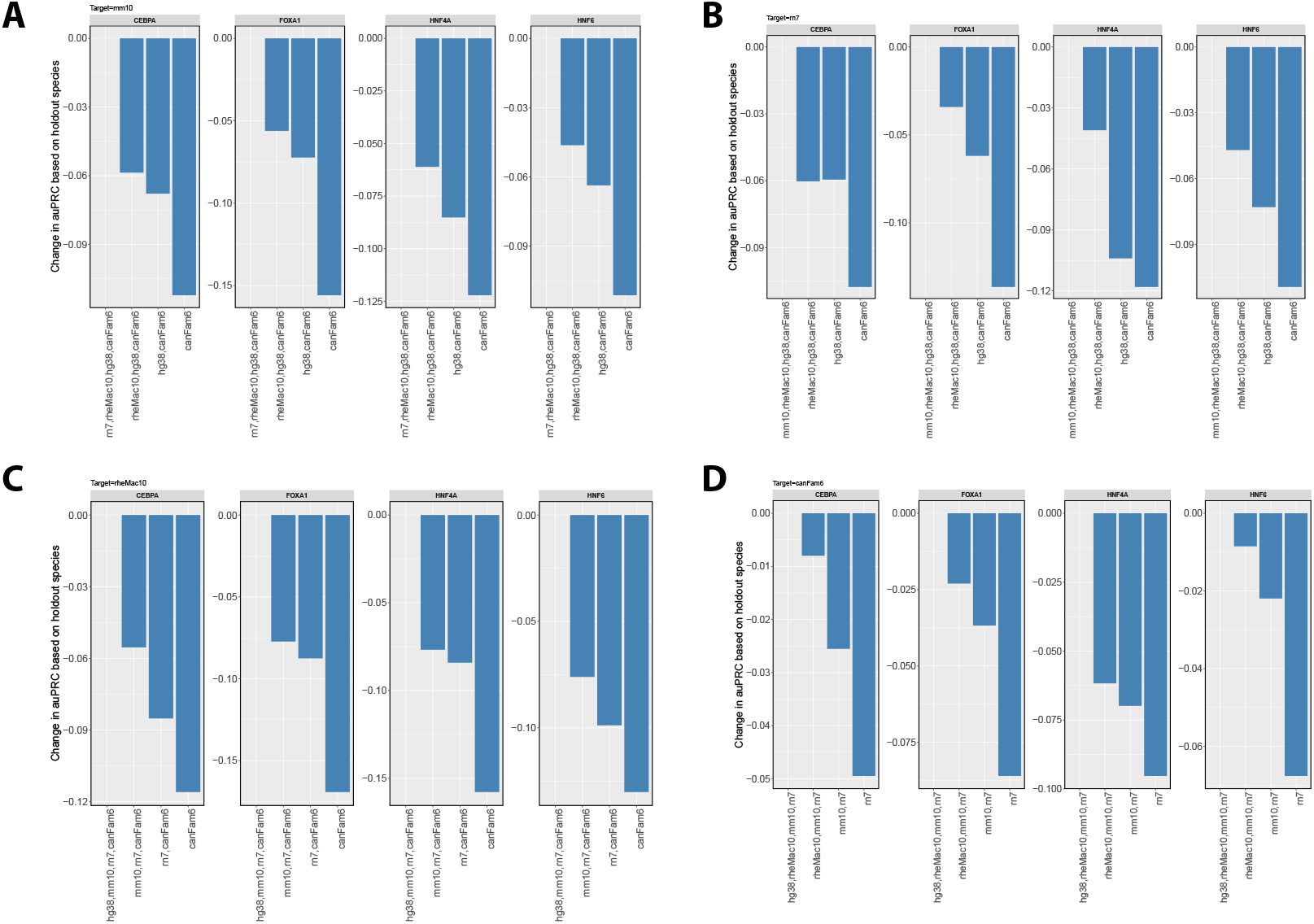
We include group holdouts under each species (other than human) as the target species.

**Figure 11:**
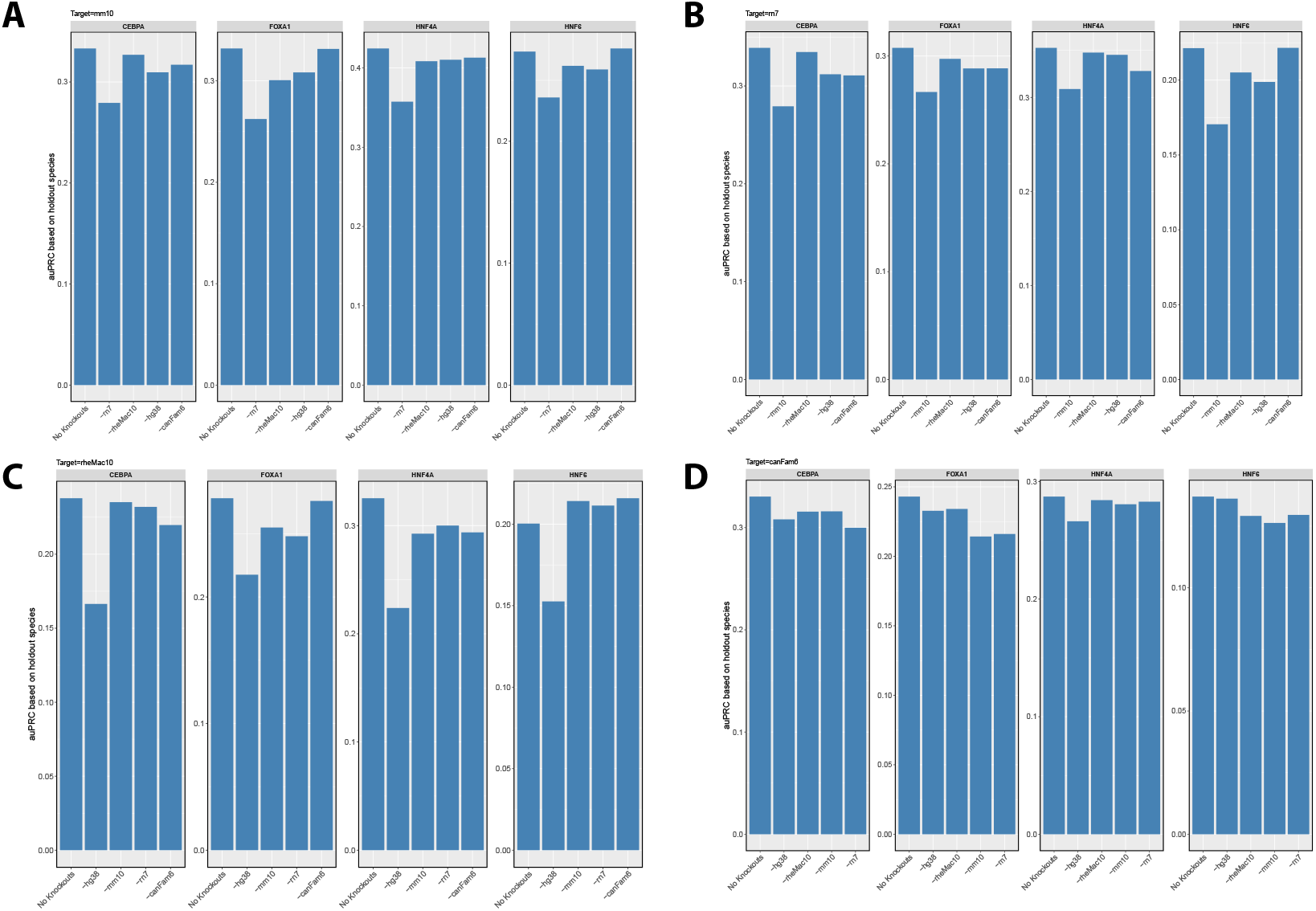
We include per species holdout under each species (other than human) as the target species.

## References

[1] Yan Hu et al. “Multiscale footprints reveal the organization of cis-regulatory elements”. In: Nature 638 (Feb. 2025), pp. 779–786. ISSN: 1476-4687. DOI: 10.1038/s41586-024-08443-4.

[2] Anusri Pampari et al. “ChromBPNet: bias factorized, base-resolution deep learning models of chromatin accessibility reveal cis-regulatory sequence syntax, transcription factor footprints and regulatory variants”. In: bioRxiv (Jan. 2025), p. 2024.12.25.630221. eprint: 2024.12.25.630221. URL: 10.1101/2024.12.25.630221.

[3] Garyk Brixi et al. “Genome modeling and design across all domains of life with Evo 2”. In: bioRxiv (2025). DOI: 10.1101/2025.02.18.638918. eprint: https://www.biorxiv.org/content/early/2025/02/21/2025.02.18.638918.full.pdf. URL: https://www.biorxiv.org/content/early/2025/02/21/2025.02.18.638918.

[4] Aman Patel et al. “DART-Eval: A Comprehensive DNA Language Model Evaluation Benchmark on Regulatory DNA”. In: arXiv (Dec. 2024). DOI: 10.48550/arXiv.2412.05430. eprint: 2412.05430.

[5] Kathleen M. Chen et al. “A sequence-based global map of regulatory activity for deciphering human genetics”. In: Nat. Genet. 54 (July 2022), pp. 940–949. ISSN: 1546-1718. DOI: 10.1038/s41588-022-01102-2.

[6] Žiga Avsec et al. “Base-resolution models of transcription-factor binding reveal soft motif syntax”. In:Nat. Genet. 53 (Mar. 2021), pp. 354–366. ISSN: 1546-1718. DOI: 10.1038/s41588-021-00782-6.

[7] Kaelan J. Brennan et al. “Chromatin accessibility in the Drosophila embryo is determined by transcription factor pioneering and enhancer activation”. In: Dev. Cell 58.19 (Oct. 2023), 1898–1916.e9. ISSN: 1534-5807. DOI: 10.1016/j.devcel.2023.07.007.

[8] Dominic Schmidt et al. “Five-Vertebrate ChIP-seq Reveals the Evolutionary Dynamics of Transcription Factor Binding”. In: Science 328.5981 (Apr. 2010), pp. 1036–1040. ISSN: 0036-8075. DOI: 10.1126/science.1186176.

[9] David R. Kelley. “Cross-species regulatory sequence activity prediction”. In: PLoS Comput. Biol. 16.7 (July 2020), e1008050. ISSN: 1553-7358. DOI: 10.1371/journal.pcbi.1008050.

[10] Žiga Avsec et al. “Effective gene expression prediction from sequence by integrating long-range interactions”. In: Nat. Methods 18 (Oct. 2021), pp. 1196–1203. ISSN: 1548-7105. DOI: 10.1038/s41592-021-01252-x.

[11] Hugo Dalla-Torre et al. “Nucleotide Transformer: building and evaluating robust foundation models for human genomics”. In: Nat. Methods (Nov. 2024), pp. 1–11. ISSN: 1548-7105. DOI: 10.1038/s41592-024-02523-z.

[12] Ling Chen, Alexandra E. Fish, and John A. Capra. “Prediction of gene regulatory enhancers across species reveals evolutionarily conserved sequence properties”. In: PLoS Comput. Biol. 14.10 (Oct. 2018), e1006484. ISSN: 1553-7358. DOI: 10.1371/journal.pcbi.1006484.

[13] Kelly Cochran et al. “Domain-adaptive neural networks improve cross-species prediction of transcription factor binding”. In: Genome Res. 32.3 (Jan. 2022), pp. 512–523. ISSN: 1088-9051. DOI: 10.1101/gr.275394.121.

[14] Baochen Sun, Jiashi Feng, and Kate Saenko. “Correlation Alignment for Unsupervised Domain Adaptation”. In: arXiv (Dec. 2016). DOI: 10.48550/arXiv.1612.01939. eprint: 1612.01939.

[15] Baochen Sun and Kate Saenko. “Deep CORAL: Correlation Alignment for Deep Domain Adaptation”. In: arXiv (July 2016). DOI: 10.48550/arXiv.1607.01719. eprint: 1607.01719.

[16] Haley M. Amemiya, Anshul Kundaje, and Alan P. Boyle. “The ENCODE Blacklist: Identification of Problematic Regions of the Genome”. In: Sci. Rep. 9.9354 (June 2019), pp. 1–5. ISSN: 2045-2322. DOI: 10.1038/s41598-019-45839-z.

[17] Ben Langmead and Steven L. Salzberg. “Fast gapped-read alignment with Bowtie 2”. In: Nat. Methods 9 (Apr. 2012), pp. 357–359. ISSN: 1548-7105. DOI: 10.1038/nmeth.1923.

[18] Shaun Mahony et al. “An Integrated Model of Multiple-Condition ChIP-Seq Data Reveals Predeterminants of Cdx2 Binding”. In: PLoS Comput. Biol. 10.3 (Mar. 2014), e1003501. ISSN: 1553-7358. DOI: 10.1371/journal.pcbi.1003501.

[19] Benoit Ballester et al. “Multi-species, multi-transcription factor binding highlights conserved control of tissue-specific biological pathways”. In: eLife (Oct. 2014). DOI: 10.7554/eLife.02626.

[20] Petr Danecek et al. “Twelve years of SAMtools and BCFtools”. In: GigaScience 10.2 (Feb. 2021). giab008. ISSN: 2047-217X. DOI: 10.1093/gigascience/giab008. eprint: https://academic.oup.com/gigascience/article-pdf/10/2/giab008/36332246/giab008.pdf. URL: https://doi.org/10.1093/gigascience/giab008.

[21] Artem Tarasov et al. “Sambamba: fast processing of NGS alignment formats”. In: Bioinformatics 31.12 (2015), pp. 2032–2034. DOI: 10.1093/bioinformatics/btv098. URL: +%20http://dx.doi.org/10.1093/bioinformatics/btv098.

[22] Siebren Frölich et al. “genomepy: genes and genomes at your fingertips”. In: Bioinformatics 39.3 (Mar. 2023), btad119. ISSN: 1367-4811. DOI: 10.1093/bioinformatics/btad119.

[23] Qianqian Liang et al. “Disease-specific prioritization of non-coding GWAS variants based on chromatin accessibility”. In: Human Genetics and Genomics Advances 5.3 (2024), p. 100310. ISSN: 2666-2477. DOI: 10.1016/j.xhgg.2024.100310. URL: https://www.sciencedirect.com/science/article/pii/S2666247724000496.

[24] Antoine de Mathelin et al. “ADAPT: Awesome Domain Adaptation Python Toolbox”. In: arXiv preprint 2107.03049 (2021).

[25] Avantika Lal et al. “gReLU: A comprehensive framework for DNA sequence modeling and design”. In: bioRxiv (Sept. 2024), p. 2024.09.18.613778. eprint: 2024.09.18.613778. URL: 10.1101/2024.09.18.613778.

[26] Martín Abadi et al. TensorFlow: Large-Scale Machine Learning on Heterogeneous Systems. Software available from tensorflow.org. 2015. URL: https://www.tensorflow.org/.

[27] François Chollet et al. Keras. https://keras.io. 2015.

[28] F. Pedregosa et al. “Scikit-learn: Machine Learning in Python”. In: Journal of Machine Learning Research 12 (2011), pp. 2825–2830.

[29] Adam Paszke et al. “PyTorch: An Imperative Style, High-Performance Deep Learning Library”. In:arXiv (Dec. 2019). DOI: 10.48550/arXiv.1912.01703. eprint: 1912.01703.

[30] Niklas Kempynck et al. CREsted: Cis Regulatory Element Sequence Training, Explanation, and design. 2024.

[31] Avanti Shrikumar et al. “Technical Note on Transcription Factor Motif Discovery from Importance Scores (TF-MoDISco) version 0.5.6.5”. In: arXiv (Oct. 2018). DOI: 10.48550/arXiv.1811.00416. eprint: 1811.00416.

[32] Shobhit Gupta et al. “Quantifying similarity between motifs”. In: Genome Biol. 8.2 (Feb. 2007), pp. 1–9.ISSN: 1474-760X. DOI: 10.1186/gb-2007-8-2-r24.

[33] Gerardo Perez et al. “The UCSC Genome Browser database: 2025 update”. In: Nucleic Acids Res. 53.D1 (Jan. 2025), pp. 1243–1249. ISSN: 1362-4962. DOI: 10.1093/nar/gkae974. eprint: 39460617.

[34] Ivica Letunic. phyloT : a phylogenetic tree generator. [Online; accessed 19. Mar. 2025]. Mar. 2025.URL: https://phylot.biobyte.de.

[35] Yaroslav Ganin et al. “Domain-Adversarial Training of Neural Networks”. In: Journal of Machine Learning Research 17.59 (2016), pp. 1–35. URL: http://jmlr.org/papers/v17/15-239.html.

[36] Faisal Mahmood, Richard Chen, and Nicholas J. Durr. “Unsupervised Reverse Domain Adaptation for Synthetic Medical Images via Adversarial Training”. In: IEEE Trans. Med. Imaging 37.12 (June 2018), pp. 2572–2581. DOI: 10.1109/TMI.2018.2842767.

[37] Mehran Javanmardi and Tolga Tasdizen. “Domain adaptation for biomedical image segmentation using adversarial training”. In: 2018 IEEE 15th International Symposium on Biomedical Imaging (ISBI 2018). IEEE, pp. 04–07. DOI: 10.1109/ISBI.2018.8363637.

